# Functional genomic analyses highlights a shift in *Gpr17*-regulated cellular processes in oligodendrocyte progenitor cells (OPC) and underlying myelin dysregulation in the aged forebrain

**DOI:** 10.1101/2020.10.26.354746

**Authors:** Andrea D. Rivera, Francesca Pieropan, Irene Chacon De La Rocha, Davide Lecca, Maria P. Abbracchio, Kasum Azim, Arthur M Butt

## Abstract

Brain aging is characterised by a decline in neuronal function and associated cognitive deficits. There is increasing evidence that myelin disruption is an important factor that contributes to the age-related loss of brain plasticity and repair responses. In the brain, myelin is produced by oligodendrocytes, which are generated throughout life by oligodendrocyte progenitor cells (OPCs). Currently, a leading hypothesis points to aging as a major reason for the ultimate breakdown of remyelination in Multiple Sclerosis (MS). However, an incomplete understanding of the cellular and molecular processes underlying brain aging hinders the development of regenerative strategies. Here, our combined systems biology and neurobiological approach demonstrates that oligodendroglial and myelin genes are amongst the most altered in the aging mouse cortex. This was underscored by the identification of causal links between signaling pathways and their downstream transcriptional networks that define oligodendroglial disruption in aging. The results highlighted that the G-protein coupled receptor GPR17 is central to the disruption of OPC in aging and this was confirmed by genetic fate mapping and cellular analyses. Finally, we used systems biology strategies to identify therapeutic agents that rejuvenate OPC and restore myelination in age-related neuropathological contexts.

## INTRODUCTION

Aging in the brain is accompanied by a gradual decline in neuronal networking and synaptic plasticity which are needed for learning and cognitive function. Notably, neuronal numbers are largely unaltered in the aging human brain (Fabricius et al., 2013, Pelvig et al., 2008). In comparison, there is evidence of gradual losses in oligodendrocytes and myelin in aging and that these changes are key factors in cognitive decline and to decreased capacity for repair following pathology (Vanzulli et al., 2020, Bartzokis, 2004, Vallet et al., 2018). Sustaining myelin and oligodendrocytes throughout life is therefore critical and is the function of a reservoir of oligodendrocyte progenitor cells (OPCs) (Rivera and Butt, 2019, Reich et al., 2018). The underlying causes of myelin loss in aging are unresolved, but there is increasing evidence that a major factor may be the decline in OPC regenerative capacity (Rachman-Tzemah et al., 2017, Neumann et al., 2019a). Hence, unravelling the fundamental changes in the aging brain is a key strategy for developing new strategies to promote repair in neurodegenerative diseases, including Multiple Sclerosis (MS) and Alzheimer’s disease (AD).

Transcriptomic studies have become increasingly important in understanding aging processes in humans and rodents (Soreq et al., 2017, Jakel et al., 2019, Nolte et al., 2001, Rachman-Tzemah et al., 2017, Ha et al., 2012). Here, using a combined transcriptomic and neurobiology approach we have identified essential oligodendroglial genes amongst the most dysregulated in the aging mouse cortex, most notably *Gpr17,* which specifically decorates a subpopulation of OPCs that are in transition to mature myelinating oligodendrocyte (MYOL) and react rapidly to brain pathology (Kari et al., 2019). In addition, we determined specific cellular signalling pathways and transcriptional networks that characterise ageing oligodendroglia. Finally, we used novel *in silico* pharmacogenomics strategies for the identification of therapeutic agents that stimulate the transcriptional networks for driving the regeneration of OPCs following demyelination and have therapeutic potential in MS and neurodegenerative diseases.

## MATERIALS AND METHODS

### Animals and tissue

All animal studies were performed in accordance with international law (European law Dir. 2010/63/UE) and national laws (UK Animals Scientific Procedures Act, 1984; Italian law DL n. 26, 4th March 2014). All procedures were reviewed and approved by the local ethical review bodies of the Universities of Portsmouth and Milan, and appropriate UK Home office Project Licence and the Italian Ministry of Health (authorization 473–2015PR to MPA). Mice were housed in groups of 4, under a 12-hr light/12-hr dark cycle at 21°C, with food and water *ad libitum.* Wild-type mice of the background strain C57/BL10 were used for gene expression profiling and induction of demyelination (see below for further details). Adult Pdgfra-CreER^T2^:Rosa26R-YFP mice (gift from Professor William D. Richardson, UCL, UK (Rivers, Young, Rizzi, Jamen, *et al*., 2008) and Gpr17-iCreERT2xCAG-eGFP mice (Viganó et al., 2016) were respectively maintained and bred at the University of Portsmouth and University of Milan facilities; offspring were ear punched and genotyped using PCR, as previously reported (Rivers, Young, Rizzi, Jamen, *et al*., 2008)(Viganó et al., 2016), and mice of the correct phenotype and age (see below for the ages used) were injected intraperitoneally (i.p.) twice a day for 5d with tamoxifen (0.1 ml of a 10 mg/mL solution, prepared in ethanol and corn oil), to induce Cre recombination and reporter expression, and brains were examined 10 days after the last injections (see below). The experiments were designed in compliance with the ARRIVE guidelines and no mice were excluded from analyses and experimental groups contained a spread of sexes. Control groups were included in all experiments, applying randomizing procedures and double-blinded analysis when possible.

### Immunohistochemistry

For immunohistochemistry, mice were perfusion fixed intracardially under terminal anaesthesia with 4% Paraformaldehyde (PFA). Brains were then dissected free and immersion fixed in 4% PFA overnight. After fixation, tissues were washed 3 times in PBS and stored at 4°C in PBS containing 0.05% NaN3 (Sigma) until use. Coronal brain sections were cut using a vibratome (Leica) at a thickness of 60 μm and used immediately or stored at 4°C in PBS/NaN3 in 24 well plates. Sections were treated for a blocking stage of either 10-20% normal goat serum (NGS) or normal donkey serum (NDS) or 0.5% bovine serum albumin (BSA) for 1-2 h, depending on the primary antibodies to be used. Sections were washed 3 times in PBS, and incubated overnight in primary antibody diluted in blocking solution containing Triton-X (0.4%): chicken anti-YFP (1:100, Abcam), rabbit anti-NG2 (1:500, Millipore), rabbit anti-GPR17 (1:00, Cayman Labs), mouse anti-APC (1:700, Calbiochem), mouse anti-MBP (1:300, Millipore), rabbit-Olig2 (1:400, Millipore), goat anti-PDGFRα (1:400, R&D). Sections were then washed 3 times in PBS and incubated with the appropriate secondary antibody (Alexa fluor^®^-488, Alexa fluor^®^-568, Alexa fluor^®^-594, Alexa fluor^®^-647) diluted in blocking solution for 1-2 h at room temperature. Following secondary antibody incubation, tissues were washed 3 times with PBS before being mounted on glass slides using Fluoromount-G (Southern Biotech). Detection of EdU (5-ethynyl-2’-deoxyuridine) was performed as per the manufacturer’s guidelines using Click-it EdU Alexa Fluor 488 imaging kit (Invitrogen).

### Imaging and analysis

Images were captured using a Zeiss Axiovert LSM 710 VIS40S confocal microscope and maintaining the acquisition parameters constant to allow comparison between samples within the same experiment. Acquisition of images for cell counts was done with x20 objective. Images for OPC reconstruction were taken using x100 objective and capturing z-stacks formed by 80-100 single plains with an interval of 0.3 μm. Cell counts were performed in a constant field of view (FOV) of 100 μm x100 μm or 200 μm x 200 μm, depending on the area analysed, in projected flattened images from z-stacks formed by 10 or 15 *z*-single plain images with 1μm interval between them; cell density was calculated as the number of cells divided by the area of the region analysed. All data were expressed as Mean ± SEM and tested for significance using unpaired t-tests.

### RNA-seq

For gene profiling, cerebral hemispheres from 6-week old and 18-month old C57/BL10 mice were removed (n=3 mice from each age), maintaining strict RNAase-free and sterile conditions throughout. RNA was extracted and processed using a RNeasy Micro kit (Qiagen), and RNA concentration was determined using Nanodrop ND-1000 spectrophotometer, after which samples were stored at −80°C until use. Material was quantified using RiboGreen (Invitrogen) on the FLUOstar OPTIMA plate reader (BMG Labtech) and the size profile and integrity analysed on the Agilent2200 RNA ScreenTape). Input material was normalised prior to library preparation. mRNA was selected using NEBNext^®^ Poly(A) mRNA Magnetic Isolation Module (E7490S) and library preparation was completed using NEBNext^®^ Ultra Directional RNA Library Prep Kit for Illumina^®^ (E7420L, New England Biolabs), following manufacturer’s instructions. Individual libraries were quantified using Qubit, and the size profile was analysed on the 2200. Individual libraries were normalised and pooled together accordingly. The pooled library was diluted to ~10 nM for storage. The 10 nM library was denatured and further diluted prior to loading on the sequencer. Paired end sequencing was performed using a HiSeq platform at 100bp PE.

### qPCR

Total RNA from experimental samples extracted above were processed for real-time qPCR of age-related changes in *Gpr17.* Total RNA was reverse transcribed (RT) by Precision Nanoscript2 (PrimerDesign, UK), following manufacturer instructions, and real-time qPCR was performed using Precision Plus qPCR Master Mix (Primer Design, UK), following manufacturer instructions, and adding to a 20 μl of PCR reaction: 10 μl of Precision Plus qPCR Master Mix, 1 μl of Gpr17 primer, 25ng of Template and 4 μl of RNAse/DNAse free Water (Gibco, USA). Amplification was performed using a Roche Lightcycler 96 according to the manufacturer’s protocol. Data normalisation was performed using the housekeeping genes *Gapdh* and *Rpl13a* and expressed as relative gene expression using the 2 ΔΔ-CT method. Assays on all samples were performed in duplicate. Primer sequences are provided in Supplementary Table 1.

### Meta-analysis of generated datasets

Normalised datasets generated by RNA-seq analysis using edgeR using standard pipeline methods. Differential expression analysis datasets were further processed as done previously (Rivera and Butt 2019) using ConsensusPathDB, String V10.5 (Herwig et al., 2016, Szklarczyk et al., 2015) and STITCH v5.0 (Guo et al., 2007a). Normalised dataset of genes dysregulated in ageing was compared to cell specific gene database for OPC and Myelinating Oligodendrocytes (MYOL) using multiple published datasets (Cahoy et al., 2008, Lovatt et al., 2007, Zhang et al., 2014a). The top 25 most significant genes associated with aging in oligodendroglia are presented as a heatmap by ranking via FDR and relative fold change. Heatmap was constructed in R in R/Studio using the ggplot2 package.”

### SPIED/CMAP Analysis

SPIED (<u>S</u>earchable <u>P</u>latform-<u>I</u>ndependent <u>E</u>xpression <u>D</u>atabase) and CMAP (<u>C</u>onnectivity <u>MAP</u>) were used to identify small molecules predicted to induce gene signatures as the age-sensitive OPC genes (http://spied.org.uk/cgi-bin/HGNC-SPIEDXXX.cgi) as described previously (Williams, 2013). A metaanalysis was performed using cluster specific OPC genes from recent publically available single cell profiling experiment of young adult *Corpus Callosum* derived cells (https://github.com/kasumaz/AdultOLgenesis). Gene list was converted to hub genes/proteins for correlating master regulators within the totality of drug-induced genes in the SPIED database, an additional feature of the SPIED webtool (http://spied.org.uk/cgi-bin/HGNC-SPIEDXXX.cgi) (Williams, 2013). In this approach, small molecules generated recapitulate transcriptional signatures associated with stimulating young adult-associated transcriptional networks in OPCs. Gene lists were merged and CMAP interrogated for specifying pro-OPC small molecules and their target genes were assembled into a matrix in R/RStudio with the input genes as Boolean values, and correlative scores.

The R package PCAtools was used to construct a PCA plot with ggplot2 as aesthetics for colour and size grading using the CMAP output Pearson’s scorings and numbers of target genes for each drug, respectively. Target genes of LY294002 were visualised using the webtool Enrichr: https://amp.pharm.mssm.edu/Enrichr/.

### Induction of demyelination

Mice aged 6 months were deeply anaesthetised under isofluorane and a canula (Alzet, Brain infusion kit 3) was implanted at the coordinates Bregma −0.5 mm, lateral 1 mm, depth 2.5 mm, for intraventricular infusion of agents (Rachman-Tzemah et al., 2017). Three days following implantation, mice were anaesthetised under isofluorane and 2% lysolecithin (LPC, L-α-lysophosphatidylcholine; Sigma-Aldrich) in a volume of 2 μl was injected through the canula to induce demyelination in the *Corpus Callosum,* as described previously (Azim and Butt, 2011). At 5 days post injection (DPI) of LPC, the PI3K/Akt inhibitor LY294002, or sterile vehicle (DMSO in saline) in controls, was delivered into the CSF for 4 days via the implanted canula, using an osmotic minipump (10 μl/h, model 1003D; Alzet Osmotic Pumps); LY294002 was administered to provide a final concentration of 2 μM in the CSF, correcting for dilution in the ventricular volume, as described previously (Azim and Butt, 2011). To measure cell proliferation, mice were given an i.p. injection of EdU (5-ethynyl-2’-deoxyuridine; Click-it EdU Alexa Fluor 488 imaging kit, Invitrogen) at 5 and 6 days DPD at 50 mg/kg. Brains were analysed at 10 DPI for gene expression and 14 DPI for cell analysis.

### Gene expression analysis following demyelination

To determine the effects of intraventricular treatment with small molecules, compared to controls, measurement of transcripts was performed on microdissected *Corpus Callosum,* as detailed previously by the authors (Azim et al., 2017, Azim et al., 2014). In brief, maintaining strict RNAase-free and sterile conditions throughout, coronal brain sections of 500 μm thickness were cut using an adult mouse brain matrix and the *Corpus Callosum* was microdissected then flash frozen (analyses were performed on *n*=3/4 samples, each of 2 pooled *Corpus Callosum).* RNA was purified using RNA easy microkit (Qiagen), with the addition of DNAse, and cDNA was obtained using Superscript II reverse transcriptase (Invitrogen), following manufacturer’s guidelines. RNA was amplified using a kit described previously from Nugene and following manufacturers guidelines [Azim, 2017 #10726;Azim, 2014 #8656], for obtaining sufficient cDNA. Samples were run using a Roche Lyghtcycler 96 (Roche) instrument. Data normalisation was done using the housekeeping gene *Gapdh* and expressed as relative gene expression using the 2 ΔΔ-CT method [Azim, 2012 #7303]. Primers were designed by Primer Express 1.5 software and synthesized by Eurofins MG. Primer sequences are provided in Supplementary Table 1. Gene expression data are presented as Mean±SEM, and samples were compared for significance in RStudio. The pHeatmap package on R was used to construct a heatmap in RStudio using all normalised expression values in a matrix.

## RESULTS

### RNA-seq transcriptome of the aging mouse cortex

The most prominent age-related changes in the brain were explored by generating RNA-seq profiles of dissected brain cortices from 1-month old adult and 18-month aged mice (Fig. 1A-C) and further investigating to identify altered signalling and transcriptional networks using ConsensusPathDB and String V10.5 (Fig. 1D, E) (Herwig et al., 2016, Szklarczyk et al., 2015) and STITCH v5.0 (Kuhn et al., 2008). A key finding was the predominance of oligodendroglial genes amongst the most significantly altered genes in the whole brain (Fig. 1C). The most temporally regulated gene was *Gpr17,* which in the brain is expressed exclusively in oligodendroglial cells, specifically in an intermediate stage between OPCs and terminally differentiated myelinating oligodendrocytes (MYOL) (Viganó et al., 2016). This defined stage has been coined as newly formed oligodendrocytes (NFO) based on their transcriptomic profile (Supplementary Figure 1) (Zhang et al., 2014b). In addition, the highest ranked genes altered in aging were the major myelin-related genes, most notable were *Mog, Plp1, Cnp* and *Ugt8a,* as well as the less well known myelin proteins *Cldn11* (Bronstein et al., 2000), Tspan2 (Yaseen et al., 2017), and *Mal,* which regulates recruitment of PLP to myelin (Bijlard et al., 2016). These trends were verified by Gene Ontology (GO) analysis, which identified the main biological processes as those associated with ECM organization and gliogenesis/differentiation, and specifically oligodendrocyte differentiation and myelination (Fig. 1D). String V.10.5 Network Visualisation revealed that the most transcriptionally reshaped landscapes in the aging cortex were associated with the control of cell cycle, and protein sub-networks coupled to ECM remodelling and myelination, together with a transcriptional subnetworks associated with *Gpr17* (Fig. 1E). The ECM plays a pivotal role in oligodendrocyte differentiation (Lourenço and Grãos, 2016) and increased stiffness of the ECM is related to age-related deterioration of OPC function (Nolte et al., 2001). Overall, these unbiased statistical analyses signify oligodendroglial genes as highly susceptible to age-related changes in the mouse cortex.

**Fig.1:**
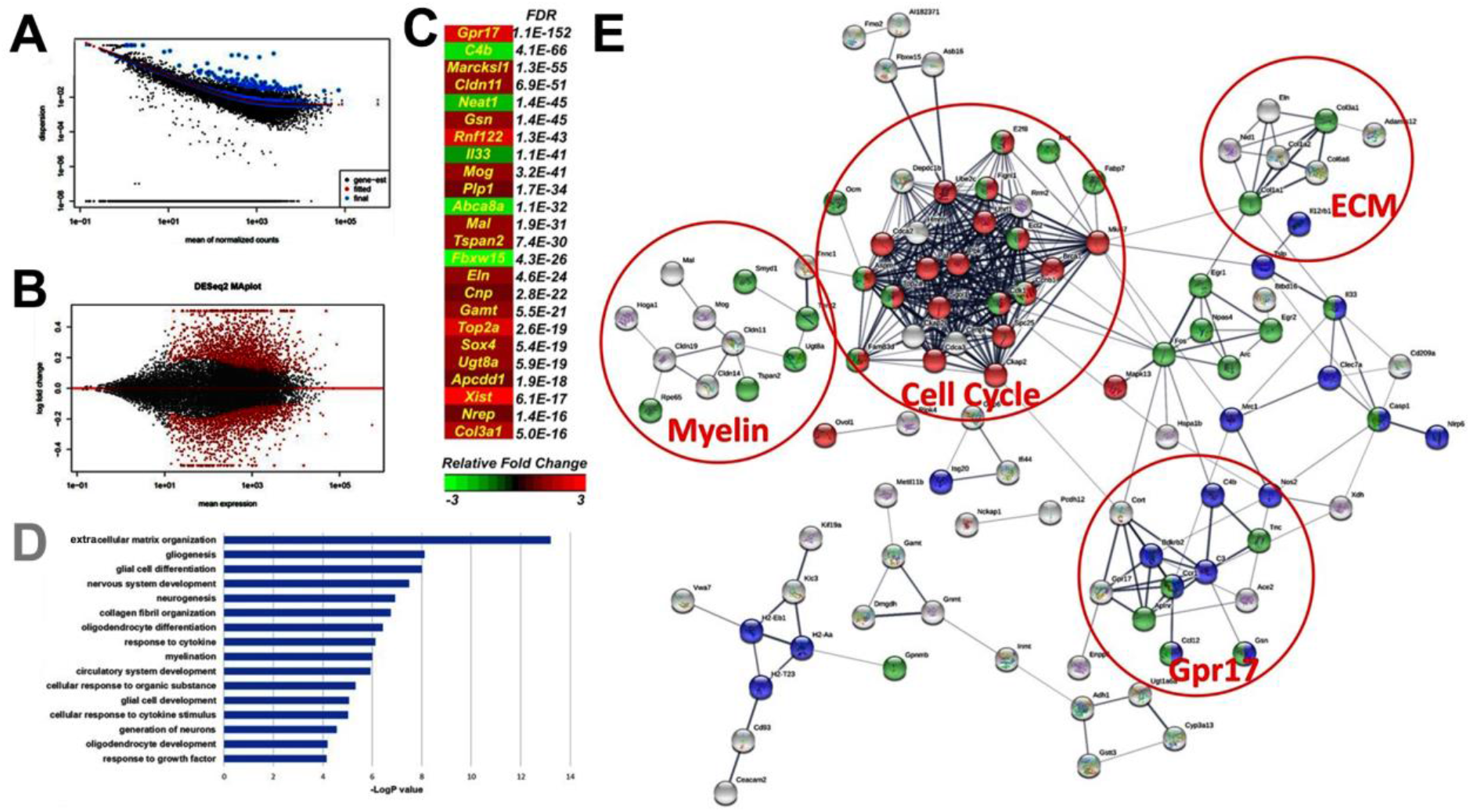
Transcriptomic characterisation of the ageing-inducing genes in the brain. (A) QC of Datasets, analysis and dispersion plot of normalised mean gene counts followed expected trend. (B) MA plot illustrating the differential expression analysis and identified 1706 genes significantly altered between the two groups (FDR < 0.01 or pADJ < 0.01) using Deseq2 (V.1.4.2). (C) Heatmap of the most altered genes in the aging cortex ranked by FDR values and colour intensity relative to fold change. (D). Major aging-induced gene changes (threshold genes at FDR < 0.05) represented by GO analysis revealing Extracellular Matrix Organisation, Gliogenesis, Neurogenesis and Myelination among the most altered Biological Pathways. (E). Network analysis of predicted protein-protein interaction performed with STRING (V.10.5) identified an alteration of the major processes and highlighting Cell cycle (Red, FDR< 0.0127), Cell Differentiation (Green, FDR<0.0307) and Inflammatory Response (Blue, FDR<0.0243, PPI Enrichment p-Value < 1.0e-16).

### Aged OPC transcriptional signature and myelination transcriptional networks

To provide insight into the stage-specific transcriptional signatures of ageing OPC (Fig. 2) and MYOL (Supplementary Figure 1), we performed a meta-analysis of our RNA-seq database against published datasets (Cahoy et al., 2008, Lovatt et al., 2007, Zhang et al., 2014a). The results confirmed the most altered processes in ageing MYOL were associated with myelination (SFig. 1B), and at the core was *Egfr* (epidermal growth factor receptor) (SFig. 1C), which has recognised importance in oligodendrocyte regeneration and myelin repair (Aguirre et al., 2007); interestingly, our analysis implicates dysregulation of a novel *Egfr-Vinculin-Gelsolin-Cldn11* axis in the age-related changes in myelination, whereby the mechanosensitive function of EGFR is transduced by *Vcl* (Vinculin) and *Gsn* (Gelsolin) which, with *Cldn11* (claudin-11), regulate the anchoring of the actin cytoskeleton to the ECM through integrins, which is essential for myelination (Bronstein et al., 2000, Zuchero et al., 2015). In aging OPCs, GO analysis demonstrated the highest ranked shifts in the cellular machinery were related to neural cell development, negative regulation of cell signalling and ECM organization (Fig. 2B). STRING analysis uncovered the key aging OPC gene networks with the largest transcriptomic hub were related to the cell cycle operating downstream of signalling via the ECM and a *Pdgfra-Gpr17* axis (Fig. 2C). Further exploration of age-induced OPC gene networks unravelled *Gpr17* as a multifactorial regulator central to numerous pro-oligodendroglial mechanisms, in addition to its receptor function (uracil nucleotides and cysteinyl leukotrienes) during the transition between OPC and MYOL (Chen et al., 2009). Our analysis identified novel interactions between *Gpr17* and OPC differentiation, synaptic signalling and the ECM, together with prominent interactions between *Gpr17* and other G-protein couple receptors, including *P2yr12* which mediates OPC-ECM interactions that regulate differentiation (Dennis et al., 2012). *Gpr17* is an upstream hub for genes that encode for synaptic proteins in OPCs (Fig. 2D), via the cell-adhesion protein *Dchs1* (Dachsous Cadherin-Related 1) and *Rasgrf1* (Ras Protein Specific Guanine Nucleotide Releasing Factor 1), which play essential roles in synaptic plasticity (Miller et al., 2013, Seong et al., 2015), and in the OPC gene network interconnect *Gpr17* with *Cacng4* (or Stargazin), together with the synaptic proteins *Shank3, Homer2, Nrxn1/2* and *Nlgn3,* which regulate synaptic targeting of AMPA receptors and bi-directional stabilization of the pre- and post-synaptic membranes (Chen et al., 2000, Dean and Dresbach, 2006, Shiraishi-Yamaguchi and Furuichi, 2007). Notably, Stargazin targets AMPA receptors to the OPC cell membrane (Zonouzi et al., 2011), and AMPA receptors regulate OPC proliferation, differentiation and myelination (Larson et al., 2016). The aged OPC transcriptional signature locates *Gpr17* at the core of these OPC signalling networks that are most altered in the aging brain.

**Fig. 2:**
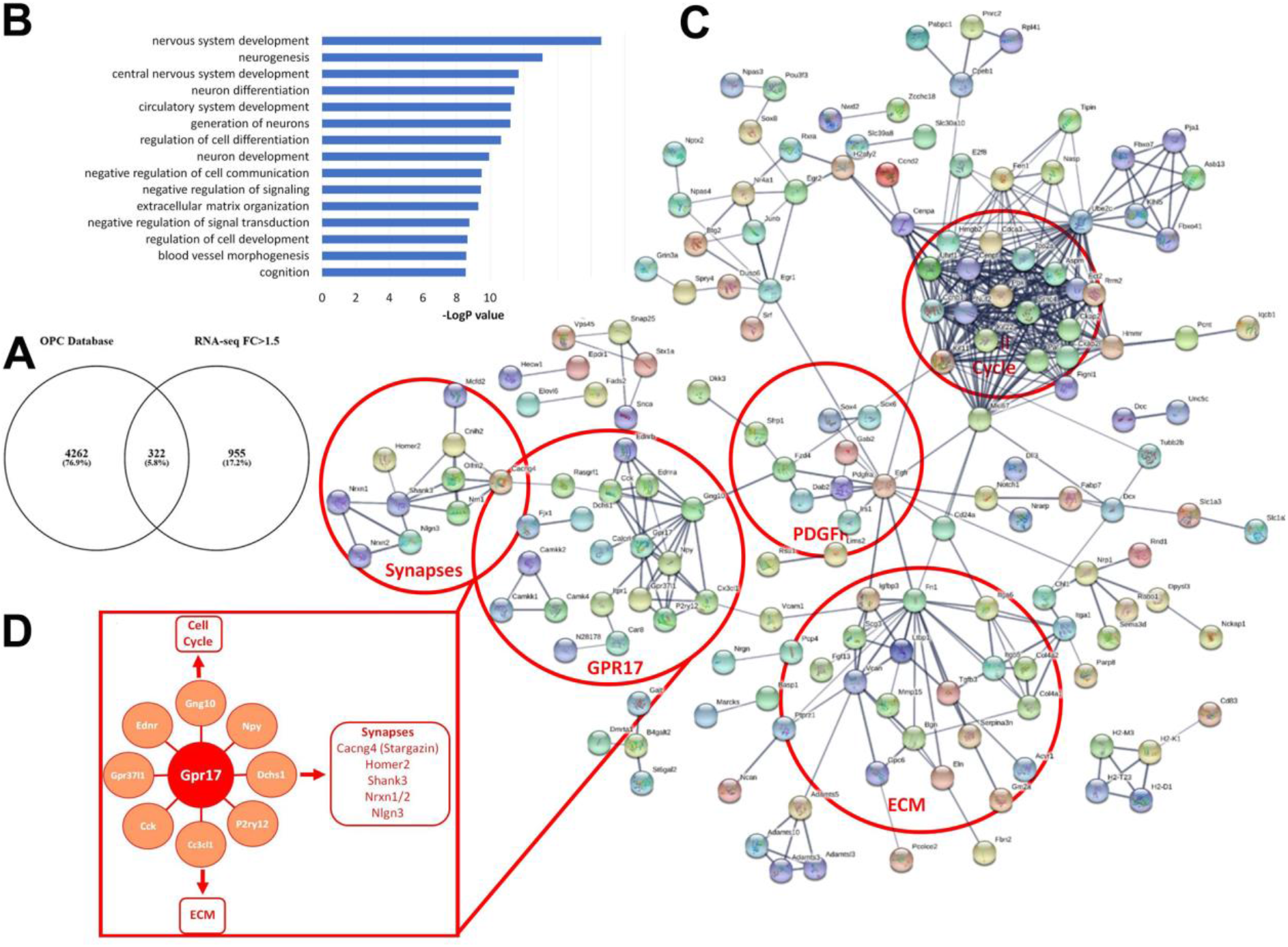
Age-related transcriptional network alterations in OPCs. (A) Meta-analysis of RNA-seq database of OPCs identified 322 core genes. (B) GO analysis identified disruption in OPC Biological Pathways related to Neural Cell Differentiation, Negative Tegulation of Cell Communication, Cell Signalling, Signal Transduction and ECM Organization. (C) STRING analysis elucidates the predicted interactions of OPC genes altered in ageing (PPI Enrichment p-Value <1.0e-16). The circles represent groups of genes active along common pathways. (D) Notably, Cell cycle genes are associated through Mki67 to the Pdgfra/Egfr and ECM nodes. Moreover, highlighted are the Gpr17 (and Synapses nodes), and are downstream targets of Pdgfra through Fzd4.

### Dysregulation of GPR17 and oligodendrocyte differentiation in aging

The transcriptomic data implicate *Gpr17* and changes in OPC regulatory networks as central to the ageing process. To investigate how these are translated into cellular changes, we examined substages of the OL lineage in the *Corpus Callosum* in Pdgfra-CreERT2:Rosa26R-YFP and Gpr17-iCreERT2xCAG-eGFP mice (Fig. 3). First, Pdgfra-CreERT2:Rosa26R-YFP mice aged 3- and 18-months were injected with tamoxifen twice a day for 5 days to induce YFP expression in OPCs. After 10 days following genetic recombination, immunostaining was performed for NG2, GPR17 and APC to identify the key stages between OPCs and terminally differentiated MYOL (Fig. 3A).

**Fig.3:**
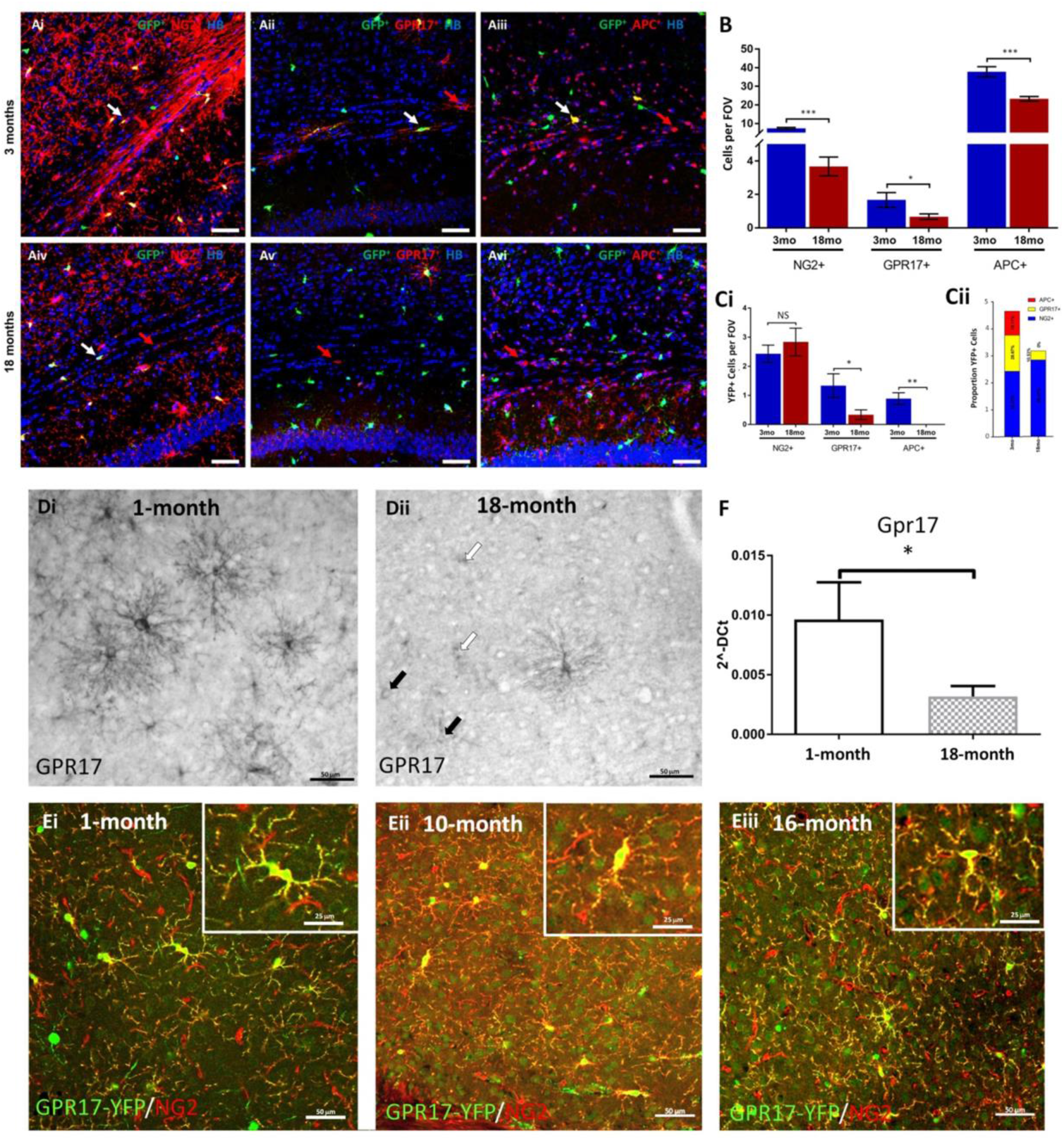
Dysregulation of Gpr17 and Oligodendrocyte differentiation. (A) Immunostaining of Pdgfra-CreERT2:Rosa26R-YFP mice aged 3-(top panels) and 18-months (lower panels) after 10 days following genetic recombination, demonstrating a reduction in the number of NG2+ OPCs (Ai, Aiv), GPR17+ OPCs/NFOs (Aii, Av), and APC+ terminally differentiated MYOLs (Aiii, Avi). Scale bar sizes are given in the panels. (B) Quantification of cell numbers per FOV showing dramatic disruption of OPCs and later stage OLs from 3 to 18 months. (C) YFP+ cell quantification in defined differentiation stages relative to total number of cells illustrating a marked decline in OPC differentiation into GPR17+ OPC/NFOLs and a complete loss of cells differentiating in to APC+ over this period. Quantification of recombined YFP+ cells presented as cell counts in ci or as a percentage in Cii. (D) Chromogenic characterisation of GPR17 expression in 1 month old (Di) and 18 months old (Dii) cortex in wildtype C57BL/6 mice indicating a loss of GPR17+ expression in ageing. GPR17 densely decorates somata and processes in 1 month old (Di), however in 18 months old OPCs are either dimly immunostained (filled arrows) or in many cases only the cell somata are immunopositive (open arrows) (dii). (E) Temporal characterisation of GPR17+/eGFP+ in Gpr17-iCreERT2xCAG-eGFP after 10 days following recombination. Immunostaining shows a gradual age-related decline in the densities of recombined OL lineage cells. (F) qPCR quantification of Gpr17 expression in young and aged cortices (p<0.05). Data are expressed as 2-dCt (n = 3 mice per group; *p<0.05, **p<0.01, ***p<0.001). Scale bars 50 μm; Scale bars in insets 25 μm.

Cell counts indicated that at 18-months there were significant decreases in NG2+ OPCs and GPR17+ OPCs/NFOs, together with a significant decrease in APC+ MYOL (Fig. 3A, B). Quantification of Pdgra-YFP+ cells in each differentiation stage relative to total number of cells clearly shows the marked decline in differentiation of Pdgra-YFP+ OPC into GPR17+ OPC/NFO and a complete absence of subsequent differentiation into APC+ MYOL over the two week experimental period (Fig. 3C). Thus, fate-mapping reveals that differentiation of aged OPC is disrupted at the intermediate GPR17+ phase that regulates the transition from OPC to MYOL (Viganó et al., 2016). To examine this further, we used chromogenic immunostaining (Fig. 3D), inducible expression of eGFP in Gpr17+ cells (Fig. 3E) and qPCR, which confirmed that Gpr17 mRNA is significantly and markedly decreased in aging (Fig. 3F). Chromogenic immunolabeling is exceptionally sensitive and demonstrates that in the aging brain, few cells exhibit the dense immunostaining of cellular processes that is characteristic of younger brain (Fig. 3Di), and instead OPCs are either dimly immunostained (Fig. 3Dii, filled arrows) or in many cases the cell somata alone are immunopositive (Fig. 3Dii, open arrows). Chromogenic labelling suggests GPR17+ cells persist in the aging brain, but that GPR17 expression is markedly reduced, and this was confirmed using tamoxifen inducible Gpr17-eGFP mice (Fig. 3E). Overall, the results indicate that dysregulation of GPR17 is central to age-related disruption of OPCs and their differentiation into MYOL.

### Identification of small molecules that rejuvenate ageing OPCs

Both the transcriptomic and fate-mapping/immunohistochemical findings demonstrate that disruption of OPCs and MYOL are major factors in the aging brain. We therefore used the SPIED/CMAP database to identify small molecules that recapitulate transcriptional changes in younger OPCs (Fig. 4), as previously described (Rachman-Tzemah et al., 2017). We interrogated the gene sets for young adult and aged OPCs generated above, together with previously curated single cell RNA-sequencing gene sets of young adult *Corpus Callosum.*

**Fig. 4:**
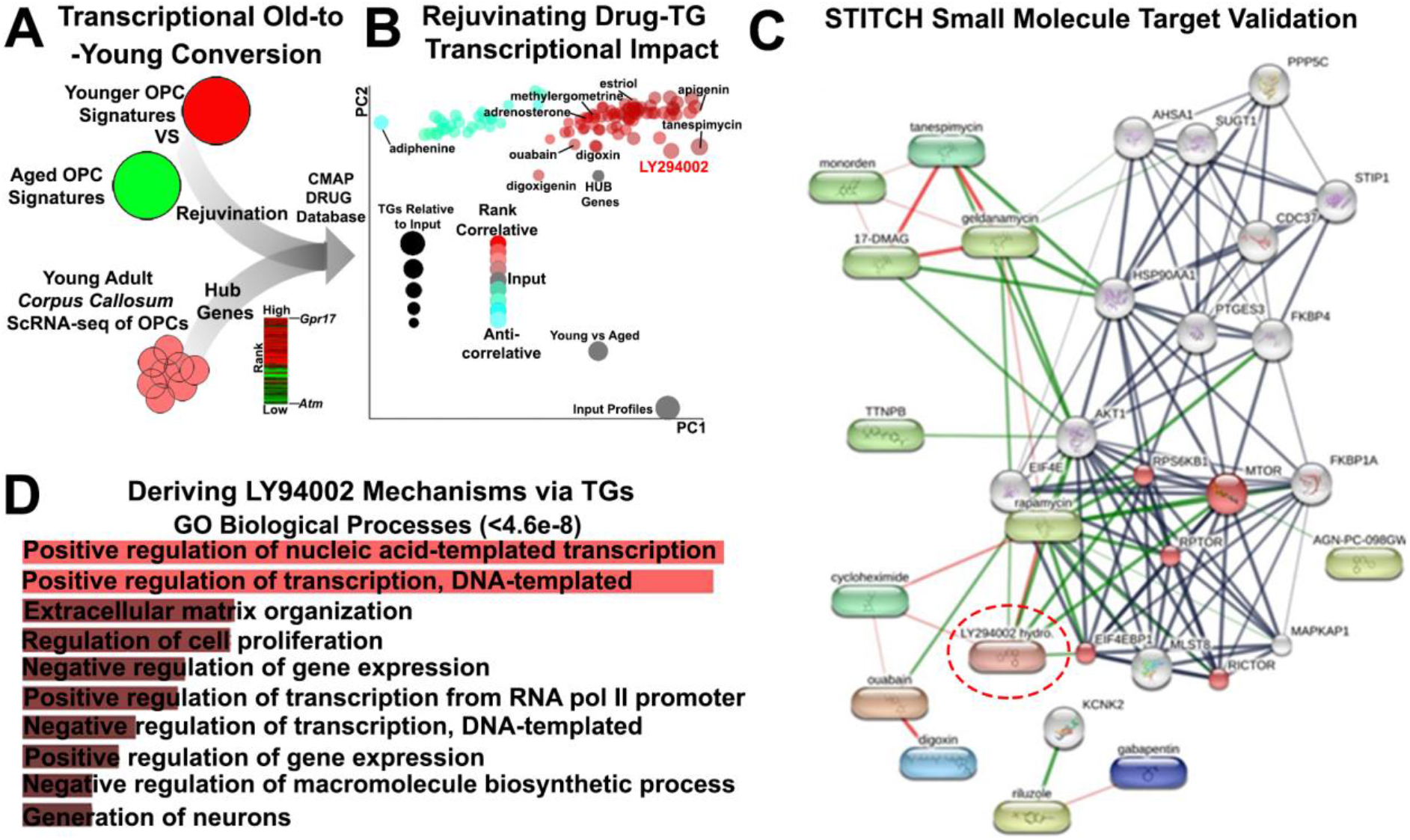
Pharmacogenomic identification of therapeutic agents for rejuvenating OPCs in a mouse model of demyelination. (A) Overview of the meta-analyses performed for assembling transcriptional signatures using datasets generated in the present study and single cell datasets of young OPCs for the detection of master regulators in the data. Combined gene lists interrogated via the CMAP database. (B) Visualisation of drugs in a principle component plot by their target genes (TGs) reflected in the size of points and coloured using correlation scores. (C) STITCH protein target analysis of pharmacogenomic drugs predicted to rejuvenate OPCs identifies LY294002 as working through the mTOR pathway (Red, FDR<9.3e-06) (PPI enrichment p-Value 3.11e-16). (D) Biological Processes of LY294002 target genes (TGs) using the Enrichr tool.

In this strategy, young adult OPC-core genes were transformed to co-expression hub genes (against drug connectivity mapping databases (Williams et al., 2013), that highlight master regulators and *Gpr17* was the most highly correlated hub (Figure 4A), which fully validates the genomic and neurobiological data presented above. We then used two distinct approaches to identify small molecules that have the potential to rejuvenate aging OPCs, by interrogating the core OPC genes across the entire SPIED/CMAP database, presented in figure 4B as a dimensionality reduction plot, in addition to a STRING chemical-protein target analysis of all OPC genes differentially expressed in the aging brain (Fig. 4C). Significantly, these two separate approaches identified the same small molecules with the potential to specifically rejuvenate ‘stemness’ in ageing OPCs (Fig. 4B, C), and none of these small molecules were predicted to act on MYOL (Supplementary Figure 2). In OPCs, a number of cardiac glycosides (digoxin, digoxigenin and ouabain) were highlighted as having OPC rejuvenation potential by regulation of mTOR signalling (Figure 4B, C). Notably, the small molecule with the highest number of target genes was LY294402, a known modulator of PI3K-Akt-mTOR signalling, which is at the centre of the OPC rejuvenating drug network (Fig. 4B, C), and is known to regulate OPC differentiation and myelination (Ishii et al., 2019). Analysis of the biological processes of LY-294002 target genes (TGs) in OPCs using Enrichr identified positive regulation of transcription, as well as ECM interactions and regulation of cell proliferation, as key mechanisms of action LY294002 in OPCs (Fig. 4D). Overall, these analyses highlighted LY294002 has a potential strategy for rejuvenating OPCs in the aging brain.

### The small molecule LY294002 identified by pharmacogenomics promotes remyelination power of OPCs in older mice

To assess the effect of LY294402 on remyelination, on remyelination of the Corpus Callosum of Sox10-EGFP mice, which identifies oligodendroglial cells at all stages of differentiation, following intraventricular infusion of the demyelinating agent lysolecithin (LPC), as detailed previously (Rachman-Tzemah et al., 2017). LPC (2%), or sterile vehicle in controls, were administered by intraventricular injection in mice aged 6 months, which was selected because at this age there is a sharp decline in the rate and overall extent of remyelination, due to reduced OPC regenerative capacity, which subsequently plateaus (Crawford et al., 2016). At 5 days post injection (DPI) of LPC to induce demyelination, LY294002 was administered by osmotic pump to provide a final concentration of 2 μM in the CSF, calculating for the dilution effect in the ventricular volume (Azim and Butt, 2011); the cell proliferation marker EdU was administered at 5 and 6 days DPI to identify proliferating OPCs and newly differentiated oligodendrocytes at 10 DPI (Fig. 5A).

**Figure 5:**
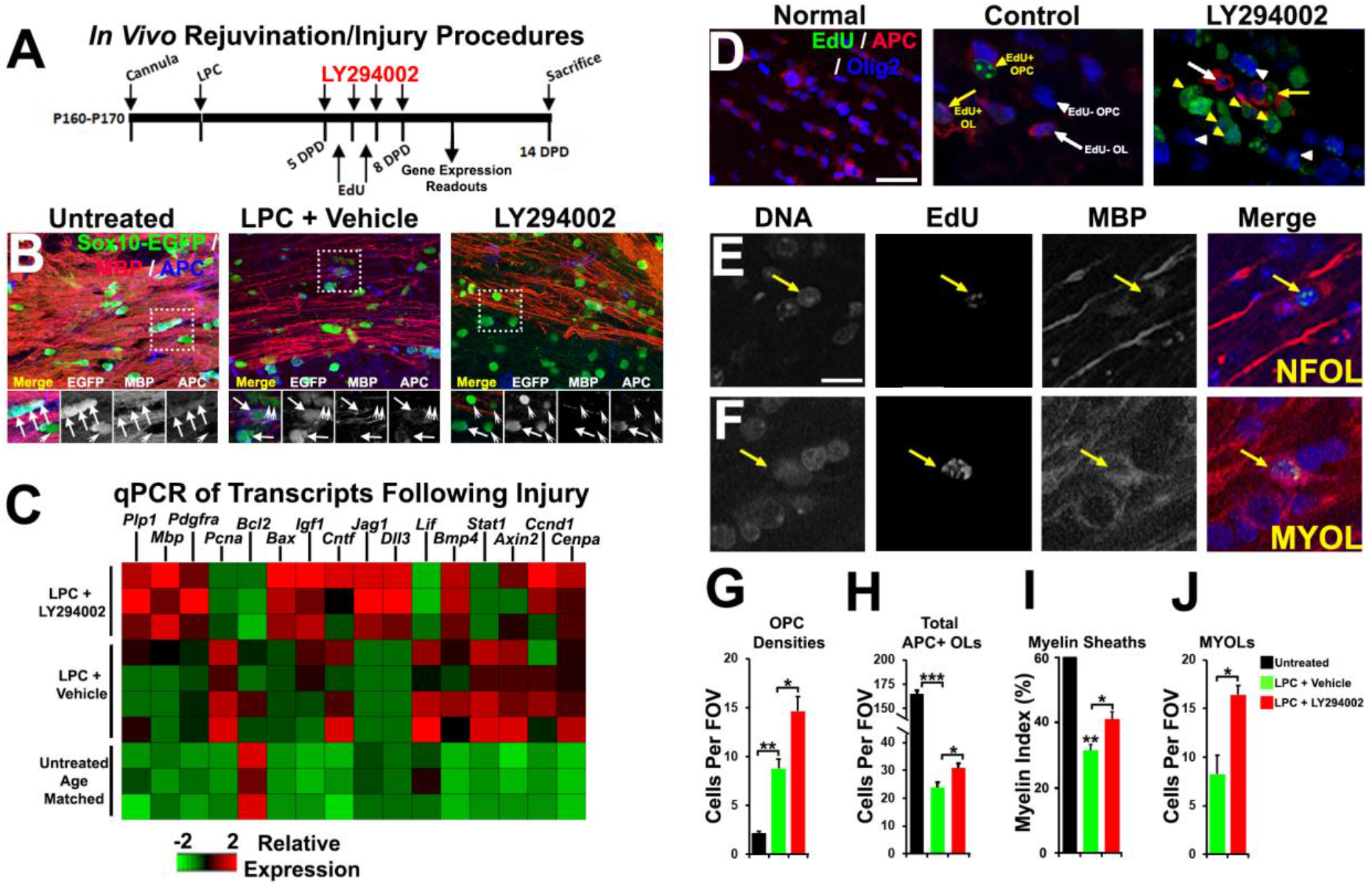
LY294002 rejuvenates OPCs in the lysolecitin model of demyelination. (A) *In vivo* demyelination procedures, EdU injections, sampling for microdissected corpus callosum tissue for qPCR and histology readouts. (B) Flattened confocal z-sections of Sox10-EGFP mice of 10 μm thickness, immunostained for myelin with MBP and differentiated OL somata via APC. (C) Heatmap of genes measured from microdissected *Corpus Callosum* tissue of untreated, lysolecithin (LPC) + control (saline/DMSO vehicle) and lysolecithin + LY294002. (D) Flattened confocal z-sections (10 μm thickness) of wildtype mice *Corpus Callosum* at 14 DPI of 10 μm thickness immunostained for Olig2 for OL lineage cells, APC and EdU for detection of cells which cycled prior. (E,F) Higher magnification of box in h to exemplify NFOLs and MYOLs quantified which incorporated EdU prior in the lineage. (g-j) Quantification of averaged Sox10-EGFP+/APC-cells and PDGFRα+ OPCs in the *Corpus Callosum* in G and total numbers of APC+ OLs in the *Corpus Callosum* in h. (G-J) Quantification of the myelin index in I and mature myelinating oligodendrocytes (MYOLs) in J. ns = no significance; * = p< 0.05; ** = p< 0.01; *** = p< 0.001. Unpaired students t-test and ANOVA posthoc applied where appropriate.

Immunolabelling for MBP confirmed evident demyelination in the Corpus Callosum at 10 DPI following LPC injection, together with an apparent decrease in the overall number of Sox10+ oligodendroglia and APC+ mature myelinating oligodenrocytes compared to controls, and there was these were evidently improved by LY294002 treatment (Fig. 5B). These effects of LY294002 were confirmed by qPCR of the microdissected Corpus Callosum of aged matched untreated mice, LCT-treated and LCT+LY294402, which demonstrates that the major OPC and oligodendrocyte transcripts *Pdgra*, *Plp1* and *MBP* were all increased significantly in LY294402 compared to LCT and controls (Fig. 5C); in addition, LY294002 had pro-oligodendroglial and anti-inflammatory effects, e.g. *Igf1* and *Bmp4* are increased, whereas *Lif* and *Stat1* are decreased compared to LCT (Fig. 5C). Finally, we analysed the changes in distinct oligodendrocyte differentiation stages by immunolabelling for the pan-oligodendroglial marker Olig2, together with EdU and APC, a marker for terminally differentiated oligodendrocytes (Fig. 5D); in this way, we identified and quantified total numbers of Olig2+/APC-OPC and Olig2+/APC+ oligodendrocytes (Fig. 5D, G, H), together with Olig2+/APC+/EdU+ newly formed oligodendrocytes and Olig2+/ Edu-/APC+ differentiated MYOL (Fig. 5E, F, I). In addition, in mature oligodendrocytes MBP immunostaining is restricted to the myelin sheaths, whereas in newly formed EdU+/APC+ oligodendrocytes it is expressed in the somata, which were prevalent in demyelinated lesions following treatment with LY294002 (Fig. 5F, arrowheads). To assess more precisely the level of remyelination, we used the myelin index which is a measure of the myelin sheaths passing through a z-plane in *the Corpus Callosum* (Fig. 5I), as previously described (Azim et al., 2012). The data demonstrate that at 10DPI, compared to controls, there was a significant increase in OPCs and decrease in mature MYOL and in the myelin index within demyelinated lesions following LCT (Fig. 5G-I), consistent with the reported loss of oligodendrocytes and recruitment of OPCs in this model. Notably, treatment with LY294002 significantly increased the numerical density of OPCs and MYOL, together with the myelin index, compared to controls and LCT treatment (Fig. 5H-I). In addition, LY294002 significantly promoted the regeneration of newly formed oligodendrocytes compared to LCT treated mice (Fig. J). The *in vivo* effects of LY294002 validate the *in silico* pharmacogenomic analysis and identifies multiple small molecules that have the potential to rejuvenate OPC stemness and promote remyelination and repair.

## DISCUSSION

Age-related changes in myelination are proposed to be a major factor in cognitive decline and the ultimate failure of remyelination and repair in MS, although the underlying mechanisms remain unclear (Bartzokis, 2004, Neumann et al., 2019c). Our differential transcriptomic analysis demonstrated that oligodendroglial genes are amongst the most significantly dysregulated in the mouse cortex in natural aging. Notably, our results highlight *Gpr17* as a major factor affected during oligodendrocyte degeneration in the aging brain. Moreover, we unravelled key transcriptional networks and signalling pathways that are central to age-related dysregulation of myelin turnover. Finally, we identified specific pro-oligodendroglial small molecules that rejuvenate OPC stemness and promote remyelination and repair. This study unravels new mechanisms in natural aging and in neurodegenerative diseases.

### Oligodendroglial transcriptomic networks are significantly altered in aging

Transcriptomic analysis identified dysregulation of multiple biological processes that are critical for normal brain function, most significantly multiple processes that are associated with dysregulation of the ECM and myelination, together with oligodenrogliogenesis. Notably, these processes are interrelated and the ECM plays a major role in the biomolecular and biomechanical regulation of OPC stemness (Nolte et al., 2001). Our data implicated an EGFR-vinculin-gelsolin-Cldn11 axis at the centre of oligodendroglial networks that are altered in aging, which provides a potential mechanism by which the mechanosensitive function of EGFR (Tschumperlin et al., 2004, Müller-Deubert et al., 2017), is transduced via vinculin and gelsolin to regulate cell/integrin/ECM interactions and control oligodendroglial cytoskeleton dynamics and cellular spreading (Hagen, 2017, Sehgal et al., 2018, Rübsam et al., 2017). Several lines of evidence show that EGFR promote oligodendrocyte regeneration and myelin repair (Aguirre et al., 2007), and our findings indicate EGFR signalling is pivotal to multiple transcriptional networks and signalling pathways that regulate age-related changes in oligodendrocytes.

### Disruption of Gpr17+ OPCs in aging

Changes in OPCs were a major hallmark of the aging brain and specifically dysregulaton of Gpr17. In the adult healthy brain, OPCs normally proliferate at very low levels and mediate rapid repair responses by reacting to injury with increased proliferation and differentiation to mature myelinating cells (Psachoulia et al., 2009). It is significant that changes related to neural cell differentiation are the top processes affected in aging OPCs, and the largest transcriptomic hub is related to the cell cycle, consistent with evidence OPC self-renewal declines with aging (Ortega et al., 2013), which underpin the striking reduction in OPC numbers observed in the 18-month brain and age-associated loss of plastic responses of OPCs and GPR17+ cells to insults and damage. Interestingly, our transcriptomic analysis of aging OPC identified a hub of genes that encode for the synaptic proteins Stargazin, Shank3, Homer2, Nrxn1/2 and Nlgn3, which are all central to glutamatergic synapses that regulate OPC proliferation and differentiation (Larson et al., 2016). Furthermore, myelination of new neuronal connections is dependent on neuronal activity and failure of OPCs to generate new oligodendrocytes retards myelination and learning ability (Ou et al., 2019, Baroti et al., 2016, Chen et al., 2000, Richardson et al., 2006, Reich et al., 2018).

### Dysregulation of GPR17 expression in aging OPC

The transcriptomic and cell biological data all point to GPR17 as being central to age-related changes in the brain. Interestingly, GPR17 is specifically expressed by a subset of OPCs that are normally quiescent but rapidly react to insults such as brain ischemia with increased proliferation and differentiation (Viganó et al., 2016), suggesting that they may serve as “reservoir” cells specifically devoted to repair purposes. Although these cells do normally fail in repairing damage due to excessive inflammatory milieu (Bonfanti et al., 2020), under “permissive conditions” (i.e., in the presence of low inflammation levels), they actually succeed in generating myelinating oligodendrocytes and ameliorating damage (Coppolino et al., 2018). In line with data showing that increased inflammation in the aged brain is associated to reduced repair abilities, our data indicate there is a marked reduction of NG2+ OPCs and myelinating oligodendrocytes at 18-months, consistent with a recent study indicating aging OPCs lose their stem cell characteristics (Neumann et al., 2019b). More importantly, our fate-mapping study shows for the first time that there is a marked decline in replenishment of GPR17+ cells from OPCs, with a subsequent loss of myelinating oligodendrocytes. Gpr17 is a receptor for damage-associated molecules such as uracil nucleotides and cysteinyl leukotrienes and regulates the transition between OPC and MYOL (Chen et al., 2009, Fumagalli et al., 2011). The balance of evidence indicates that GPR17 delays OPC differentiation, via activation of Gα_i/o_ and inhibition of cAMP-PKA (Simon et al., 2016), and desensitization by G-protein receptor kinase phosphorylation is necessary for terminal differentiation of OPC (Daniele et al., 2014). Notably, GPR17 also functions as a sensor for brain damage (Lecca et al., 2008) and antagonism of GPR17 has a rejuvenation effect in the aging brain (Marschallinger et al., 2015). Our chromogenic immunohistochemical data and analysis of GPR17 Cre-Lox mice demonstrates major disruption of GPR17 at the mRNA and protein level in aging OPCs. In addition, we identified novel interactions in Gpr17 that are altered in aging OPCs, with a prominent interaction with Gng10 (G Protein Subunit Gamma 10), which links Gpr17 to both OPC proliferation and negative regulators of OPC differentiation, namely Pdgfra, Sox 4, Sox6 and Egfr (Baroti et al., 2016, Braccioli et al., 2018, Ivkovic et al., 2008). Our data identifies a pivotal role for Gpr17 dysregulation in the aging brain and the decline in OPC capacity to regenerate oligodendrocytes.

### Pharmacogenomic screening identifies LY294002 as a therapeutic target for stimulation OPCs in the context of remyelination in older mice

We employed two separate pharmacogenomic approaches for determining: (1) the most optimal therapeutic agent for enhancing the densities of OPCs in the *Corpus Callosum* and their terminal differentiation into NFOLs; (2) Small molecules capable of reshaping aged transcriptional networks into their younger counterparts where their efficiency for myelin generation is more pronounced. Small molecules obtained in our analysis included those that target mTOR regulated cellular processes, including lipid metabolism, nucleotide synthesis and translation (Figlia et al., 2018), and the PI3K/Akt/mTOR signalling inhibitor LY294402 was identified as the most potent small molecule for shifting aged OPC into the transcriptional hallmarks that are characteristic of younger OPCs. LY294002 target genes in rejuvenating OPCs included ECM reorganisation, which we show above is dysregulated in aging OPCs. Importantly, we demonstrate that LY294003 promotes regeneration of OPCs and oligodendrocytes in vivo following demyelination induced by the toxin lysolecithin in 6-month old mice, at which age the pace of remyelination is drastically impaired compared to young adults (Kazanis et al., 2017, Crawford et al., 2016). These data validate the pharmacogenomic data and demonstrate that small molecules identified using this approach have considerable potential in reversing the decline in OPC function in aging and promoting remyelination and repair.

### Conclusions

Our unbiased transcriptomic analysis identified oligodendroglial genes amongst the most altered in the ageing mouse cortex, highlighting *Gpr17* as a major factor in the disruption of the regenerative capacity of OPCs and decline in myelination. Unravelling the key transcriptional networks and signalling pathways that are central to age-related dysregulation of OPCs, enabled us to pharmacogenomically stimulate OPC rejuvenation in the context of remyelination. This study provides a framework for future investigations in the field for targeting cellular mechanisms underlying the decline in cellular plasticity. Moreover, the usefulness in employing systems biology tools for counteracting the age-related decline in regenerative capacities and pathology.

---

**Supplementary Fig. 1:**
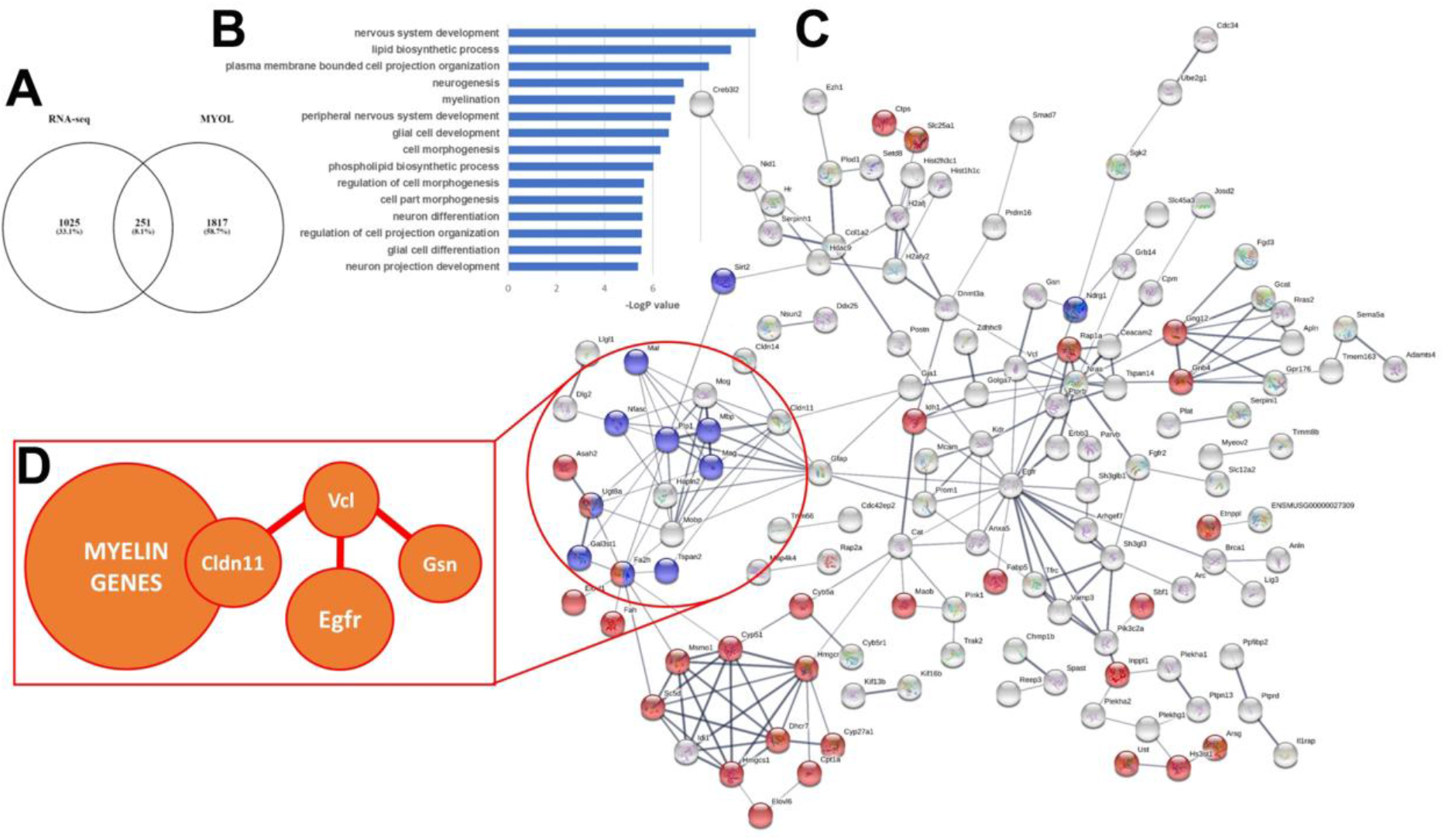
Age-related transcriptional network alterations in MYOLs. (a) Meta-analysis of RNA-seq database of MYOLs identified 251 core genes. (b) GO analysis identified the MYOL Biological Pathways mostly altered in aging including Lipid Biosynthetic processes and Myelination. (c) STRING network analysis demonstrates the predicted connectivity of MYOL genes altered in aging (PPI enrichment p-Value: 2.44e-15); The analysis identified Metabolism (Red, FDR< 3.4e-05); Myelination (Blue, FDR<5.55e-07). (d) Notably, STRING analysis identified a core network between EGFR to Myelin via Vcl, Gsn and Cldn11.

**Supplementary Fig. 2:**
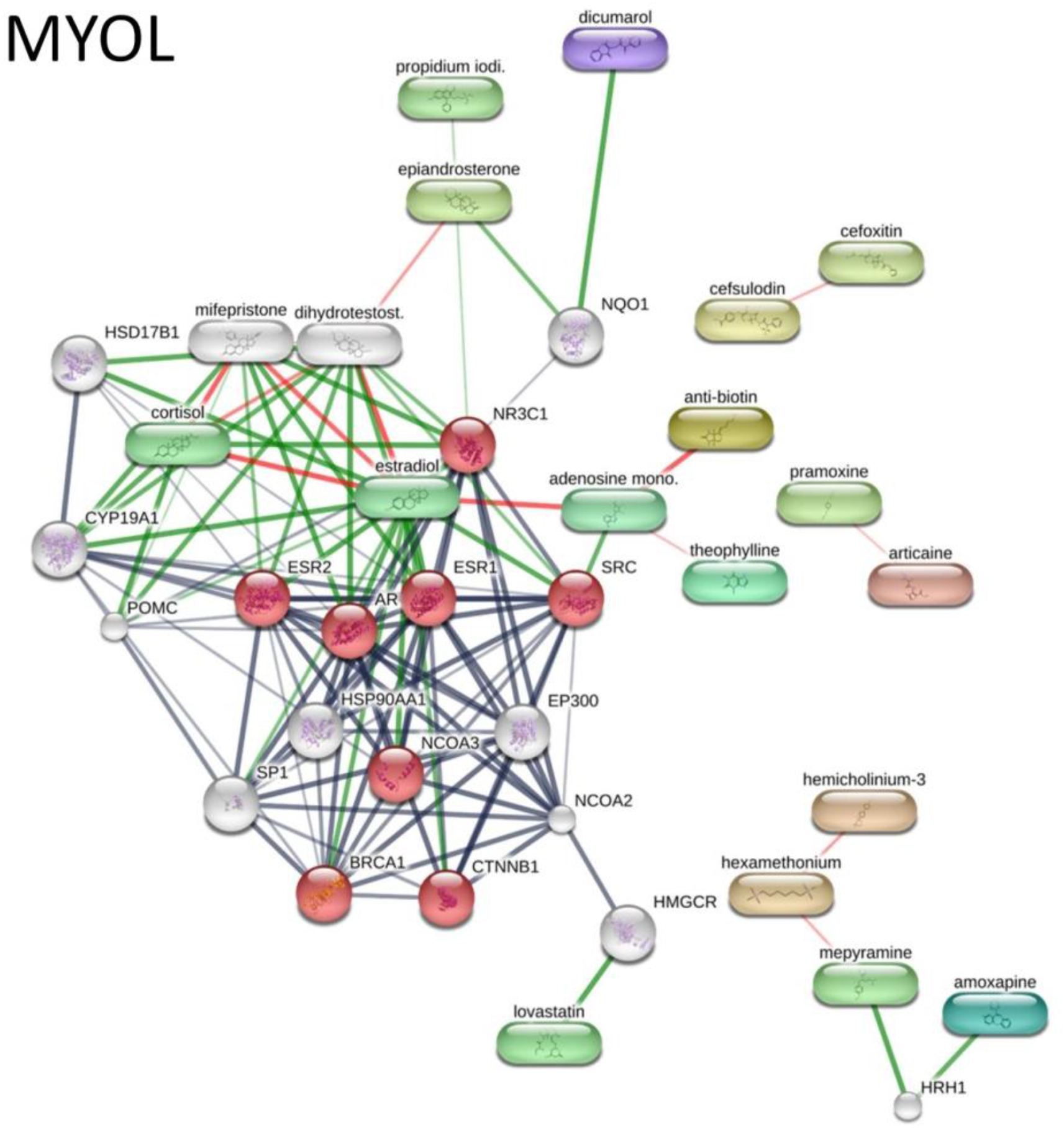
In silico drug discovery for stimulating oligodendrocyte maturation and myelination. STITCH Protein target analysis of pharmacogenomic drugs predicted to rejuvenate or MYOL (b). MYOL stimulators include Estradiol and Cortisol as targetsof the core network of Intracellular steroid hormone receptor signalling pathway (Red, FDR<1.84e-07) (PPI enrichment p-Value 2.97e-07).

**Supplementary Table 1:**
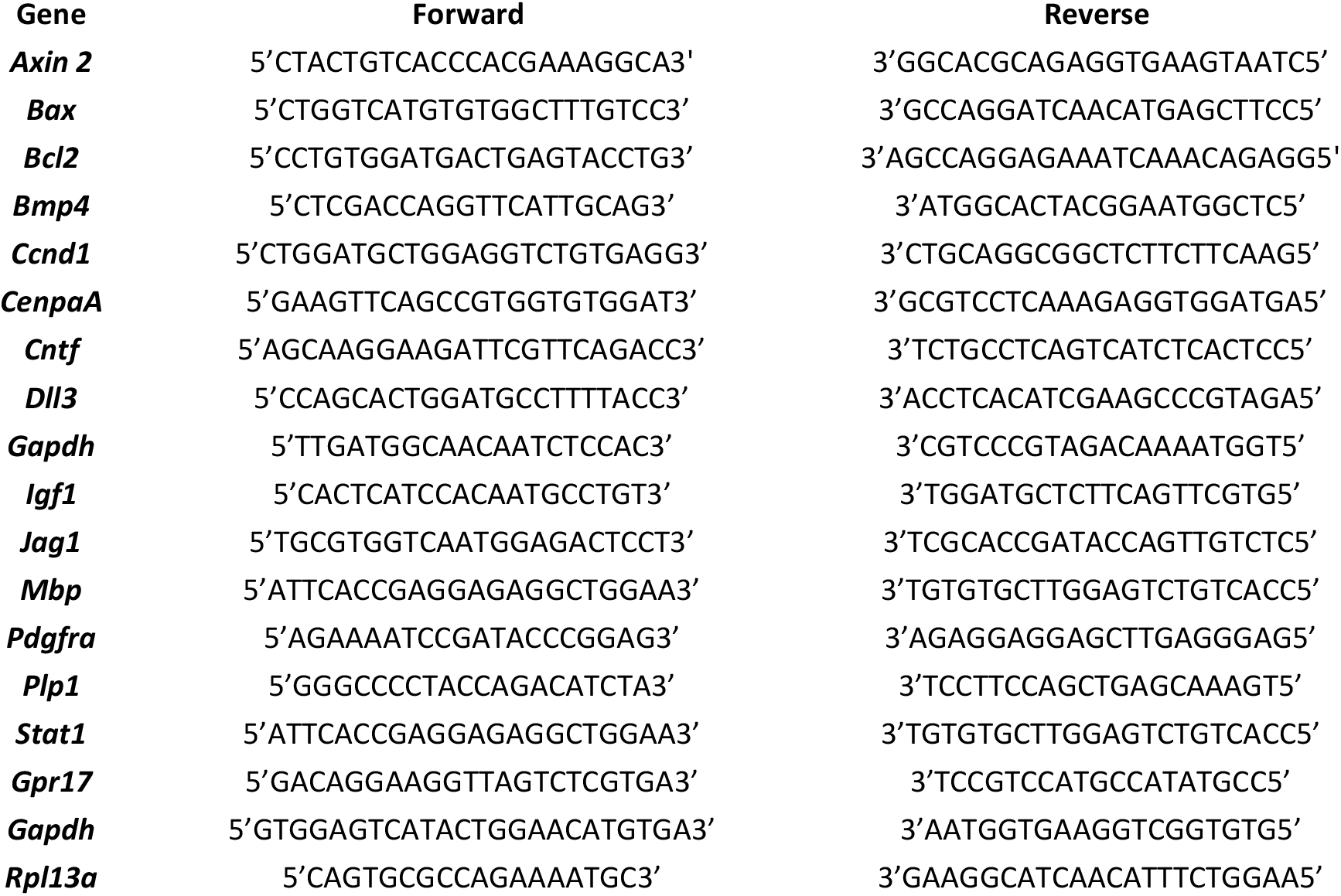
List of qPCR primer sequences used.

